# *Myo*-inositol Oxygenase Overexpression Rescues Vitamin C Deficient Arabidopsis (*vtc*) Mutants

**DOI:** 10.1101/2021.02.24.432757

**Authors:** M. Acosta-Gamboa Lucia, Nepal Nirman, Medina-Jimenez Karina, C. Campbell Zachary, S. Cunningham Shannon, Lee Jung Ae, Lorence Argelia

## Abstract

Biosynthesis of l-ascorbate (AsA) in plants is carried out by a complex metabolic network, which involves d-mannose/l-galactose, d-galacturonate, l-gulose, and *myo*-inositol as main precursors. Arabidopsis lines over-expressing enzymes in the *myo*-inositol pathway have elevated AsA, accumulate more biomass of both aerial and root tissues, and are tolerant to abiotic stresses as shown by manual and digital phenotyping. We crossed *myo*-inositol oxygenase (*MIOX4*) over-expressers with two low-vitamin C mutants (*vtc*1-1 and *vtc*2-1) encoding enzymes involved in d-mannose/l-galactose route. The purpose of developing these crosses was to test MIOX4’s ability to restore the low AsA phenotype in mutants, and to assess the contribution of individual biosynthetic pathways to abiotic stress tolerance. We used a powerful high-throughput phenotyping platform for detailed phenotypic characterization of the Arabidopsis crosses with visible, fluorescence, near-infrared and infrared sensors. We combined digital phenotyping with photosynthetic parameters and soil water potential measurements. Our results show that *MIOX4* is able to restore the AsA content of the mutants and the restored lines (vtc+MIOX4) show high AsA, enhanced growth rate, accumulate more biomass, and display healthier chlorophyll fluorescence and water content profiles compared to controls.

**Highlights:** Constitutive expression of a *myo*-inositol oxygenase restored vitamin C deficient (*vtc* mutants). The restored lines have elevated ascorbate content and are tolerant to abiotic stresses. Under normal and abiotic stress conditions, the restored lines have enhanced biomass and increased water retention.

## Introduction

Vitamin C (L-ascorbic acid, ascorbate, AsA) is one of the main antioxidants present in plants, and a regulator of growth and development. Ascorbate contributes to photosynthesis, cell division, senescence, antioxidant defense, cell wall growth, tolerance to stresses, and acts as precursor for theronic and tartaric acids (Smirnoff and Wheeler, 2000; Gallie *et al*., 2013).

Although ascorbate is an old molecule it was until 1998 when a route for its synthesis in plants was proposed, the d-mannose/l-galactose pathway (Wheeler *et al*., 1998). A second route known as the D-galacturonate pathway involving cell wall pectin as precursor was proposed next (Agius *et al*., 2003). A third pathway involving conversion of GDP-d-mannose to GDP-l-gulose was proposed also in 2003 (Wolucka and Montagu, 2003). A forth reported pathway includes the synthesis of ascorbate using *myo*-inositol as a precursor (Lorence *et al.,* 2004).

The isolation of mutants (*vtc*1-1 and *vtc*2-1) that contain low AsA content has been a useful tool to study the role of this molecule in plant growth and development. In the *vtc*1-1 line, there is a mutation in the GDP-d-mannose phosphorylase, which decreases enzyme activity up to 40%. This mutant accumulates only 25-30% of the wild-type foliar AsA (Conklin *et al*., 1996, 1999*a*). In *vtc*2-1 there is a mutation in the GDP-L-galactose phosphorylase, which results in complete loss of its activity and accumulates only 20-30% of the AsA levels (Conklin *et al*., 2000).

Cloning of different genes involved in AsA biosynthesis has enabled the production of transgenic plants with enhanced levels of this molecule. Previous studies have shown that when the MIOX4 ORF is over-expressed, there is a 2-3 fold increase in foliar AsA levels (Lorence *et al*., 2004; Tóth *et al*., 2011; Lisko *et al*., 2014).

The *myo*-inositol oxygenase enzyme is also involved in the synthesis of cell wall components (Kanter *et al*., 2005) and is required for responses to low energy conditions (Alford *et al*., 2012). Despite these characteristics, it has been proven that Arabidopsis lines over-expressing MIOX display enhanced growth, increased biomass accumulation, and tolerance to abiotic stresses (Lisko *et al*., 2013; Yactayo-Chang *et al.,* 2018; Acosta-Gamboa *et al.,* 2020). Increased auxin, enhanced photosynthesis, and increased intracellular glucose seem to be the likely mechanisms behind the enhanced biomass phenotype of the Arabidopsis MIOX over-expressers (Nepal *et al*., 2019).

Under abiotic stresses, the redox balance in plants is disturbed and reactive oxygen species (ROS) start to accumulate. If the stress applied exceeds the anti-oxidative capacity of the cell to repair, these ROS molecules can cause irreversible damage in the cell, promoting apoptosis and senescence (Veljović Jovanović *et al*., 2017; Rasool *et al*., 2018). It is when these processes are triggered, that ascorbate is considered as an essential molecule in regulating ROS level and hormonal responses to different stresses (Foyer and Noctor, 2011; Noctor *et al*., 2018).

The study of abiotic stress tolerance is crucial to understand the effects of climate change on plant plasticity and adaption to new environmental conditions. Plants subjected to water limitation as a result of increased temperature, water depletion, or excess salts could suffer irreversible physiological damage and cell death (Miller *et al*., 2010). The combination of metabolic engineering with high-throughput phenotyping could be the key to understand how AsA triggers different cell protection mechanisms under stress conditions.

Crop productivity is one of the major concerns, when challenges of climate change increase over the years reducing crop yield. Plants respond to the environment by modulating their phenotype and yield. Improving phenomic technologies is necessary to further understand the advances in genotyping to obtain robust phenotypes and improved crop output (Furbank and Tester, 2011). Image-based phenotyping methods help better understand plant adaptation under unfavorable environments. Phenomic measurements are high-throughput, non-destructive, and unbiased (Ghanem *et al*., 2015; Gehan and Kellogg, 2017). The phenomics approach typically utilizes high-resolution cameras to capture plant images ranging from visible, fluorescence, and infrared spectra in order to quantify plant architecture, chlorosis, chlorophyll fluorescence, water content, and leaf temperatures, among other traits of interest (Fahlgren *et al*., 2015*b*).

In this work we used high-throughput phenotyping technologies to document the phenome of Arabidopsis plants over-expressing MIOX4 and grown under different abiotic stresses, including water limitation, salinity, and heat. Our data shows the importance of using multiple approaches as, high throughput phenotyping, hand-held spectrometers, soil water potential measurement, seed yield, and gene expression analysis using RT-qPCR to understand the role of AsA under abiotic stress conditions.

## Materials and methods

### Seed stocks

*Arabidopsis thaliana* (Col-0, CS-70000) seeds were obtained from ABRC (The Ohio State University, Columbus, OH). A single-insertion, homozygous line over-expressing *At*MIOX4 (*At*MIOX4L21) was developed in the Lorence Laboratory as described (Yactayo-Chang, 2011; Nepal *et al*., 2019). Vitamin C deficient mutants (*vtc*1-1 and *vtc*2-1) seeds were obtained from the Conklin Laboratory.

### Plant growth conditions

Seeds were surface sterilized sequentially with 70% ethanol, 50% bleach, 0.05% Tween 20, and rinsed with sterile water before being plated on MS media (Murashige and Skoog, 1962) supplemented with 3% sucrose. Seeds were vernalized for 3 d at 4°C before being transferred to an environment controlled chamber (Conviron, Pembina, ND) at 22 ± 1°C, 65 ± 5% relative humidity, and 160-200 µmol m^-2^ s^-1^ light intensity on a long day photoperiod (16 h day:8 h night). After true leaves formed (12 d after sowing), vigorous seedlings were transferred into 3.5’’ Kord square green pots (Hummert International, MO) containing plant growing media PRO-MIX PGX (PROMIX, PA). Plants were grown to maturity under these conditions, and this process was repeated for all the generations when crosses were performed.

### Crosses

A *AtMIOX4* over-expresser was manually crossed with *vtc* mutants (*vtc*1-1 and *vtc*2-1). Reciprocal crosses were made as previously described (ABRC, 2012). We refer to crosses as follows: a homozygous cross between MIOX4 and *vtc*1-1 is called restored line 1 (RV1), while a cross between MIOX4 and *vtc*2-1 is called restored line 2 (RV2). Ascorbate measurements and PCR screens were performed until the six generation to obtain homozygous crosses.

### DNA isolation

Leaves from 5 biological crosses were used to isolate DNA at the 5.0 developmental stage (Boyes *et al*., 2001). Genomic DNA was extracted from F0 to F6 using the CTAB method (Doyle, 1991). DNA was stored at −20°C until used for analysis. Plants with negative PCR results were discarded.

### Genotype screening by PCR

The success of crosses was determined using PCR and it was performed in each generation until homozygous reciprocal crosses were obtained. Plants were screened for presence of *kanamycin resistance gene, nptII* and *MIOX4* using the primers listed in Supplementary Table 1. Gel electrophoresis was performed on a 1% agarose gel and imaged using Molecular Imager^®^ Gel Doc^™^ XR (Bio-Rad, USA).

### Ascorbic acid measurements

Leaves were collected at developmental stage 6.1 (Boyes *et al*., 2001) in the morning (9:00 am-12:00 pm) and immediately frozen in liquid nitrogen and stored at −80°C. Ascorbate was measured using an enzyme-based spectrophotometric method (Lorence *et al*., 2004). Briefly, leaves were pulverized in liquid nitrogen, and ascorbate was extracted in 6% (w/v) fresh *meta*-phosphoric acid. Reduced AsA was measured in a reaction mixture containing 950 µl of 100 mM potassium phosphate buffer pH 6.9 and 50 µl of plant extract. For *vtc* mutants, reduced AsA was measured using 900 µl of 100 mM potassium phosphate buffer (pH 6.9) and 100 µl of plant extract. After 1 min, the decrease in absorbance was recorded at 265 nm following the addition of 20 µl ascorbate oxidase (50 units/ml) (Sigma). Oxidized AsA was measured by recording the absorbance at 265 nm, before and 20 min after the addition of 1 µl 200 mM DTT to a 1 ml reaction mixture. An extinction coefficient of 14.3 mM^-1^ cm^-1^ was used for calculations. Total AsA is reported as the sum of reduced and oxidized ascorbate.

### RNA isolation

Three biological replicates of leaves were collected at developmental stage 5.0. The Purelink^TM^ RNA mini kit (Ambion, Life Technologies, USA) was used for extraction and purification of total RNA. RNA quantity and quality were assessed using an Experion instrument (Bio-Rad).

### DNA Sequencing

The *vtc*1-1 mutant (At2g39770) contains a single base mutation (T to C) at the N-terminus (second exon). This mutation leads to a Pro22Ser substitution in the active site of GDP-mannose pyrophosphorylase, resulting in a 70% decrease in ascorbate (Pavet *et al*., 2005; Mukherjee *et al*., 2010; Kerchev *et al*., 2011; Zechmann, 2011; Zhang *et al*., 2012). In the *vtc*2-1 mutant (At4g26850), there is a single base substitution (G to A) at the predicted 3′ splice site of the fifth intron; this reduces the transcript level by 80-90% and the activity of GDP-L-galactose phosphorylase activity in leaves. This mutation results in a 70-80% reduction in ascorbate levels (Müller-Moulé *et al*., 2004; Pavet *et al*., 2005; Dowdle *et al*., 2007; Kerchev *et al*., 2011).

For primer design an alignment of the CDS and mRNAs coding for Arabidopsis GDP-mannose pyrophosphorylase (At2g39770) and GDP-L-galactose phosphorylase (At4g26850) was performed using all the nucleotides and amino acid sequences to define the target regions and conserved sites. Primers were designed using Primer3 software and the NCBI database (Supplementary Table 1). An *in-silico* PCR assay was performed to confirm primer location, efficiency, orientation, and length of each amplicon. DNA and RNA were isolated from leaves of three biological replicates of *vtc*1-1 and *vtc*2-1 crosses (RV1 and RV2), including the controls, using the methodology described above. In the case of *vtc*1-1, PCR was performed, and for *vtc2-1*, RNA was converted into cDNA using the methodology described later followed by RT-PCR.

The cDNA was subjected to 30 cycles of PCR amplification (94°C for 1 min, 68°C for 30 sec and 72°C for 25 sec) in the presence of both forward and reverse primers (Table 1). DNA was amplified for 30 cycles (94°C for 1 min, 65°C for 30 sec and 72°C for 25 sec). PCR products were cleaned using the NucleoSpin^®^ Gel and PCR Clean-up (Macherey-Nagel, USA). The amplified products were cloned into pDrive using the TA Cloning Kit (Qiagen^®^, USA), following the manufacturer’s instructions. Plasmid DNA was purified using the Plasmid Miniprep kit (Qiagen^®^), following the manufacturer’s instructions. DNA was sent for sequencing to the University of Chicago Comprehensive Cancer Center DNA Sequencing and Genotyping Facility (Chicago, IL). Results were analyzed using the VecScreen software (NCBI, USA). Chromatogram were analyzed using FinchTV. Finally, the SeaView software was used to align the sequences to identify the sites of the mutations for *vtc1-1* and *vtc2-1*.

**Table 1.**
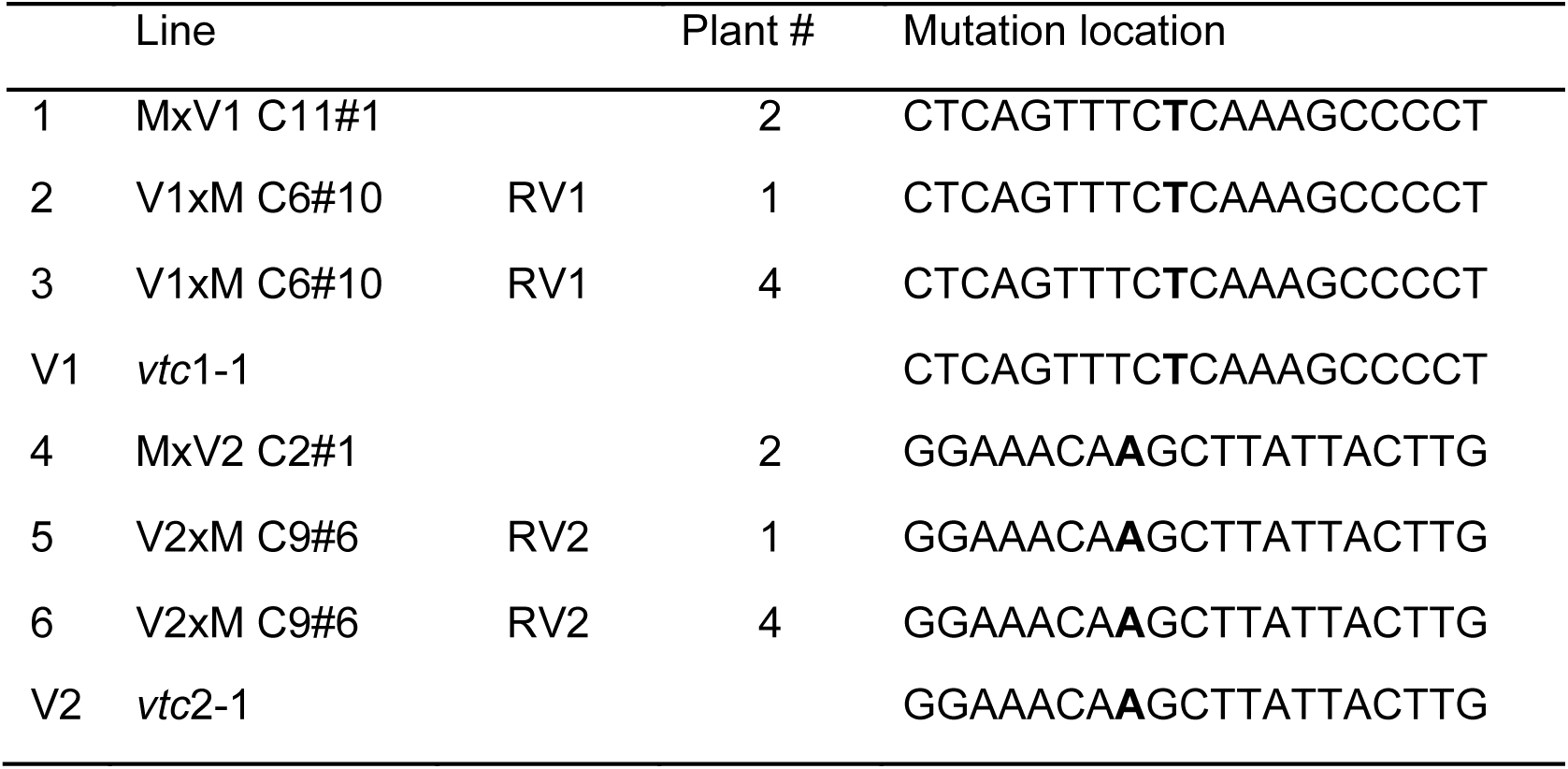
Results from DNA sequencing (n=3)

### Water limitation stress

Three days before transplanting the seedlings to soil, Quick Pot 15 RW trays were filled with dry growing media Pro-Mix PGX (Promix, PA), and their weight was recorded. Measured amounts of water were added to the dry soil and allowed to absorb for 1 h until soil reached full water saturation (100% full water capacity, FC), after which tray weights were once again recorded. After seedling establishment (approx. 20 d after germination), soil was allowed to reach four different levels of water saturation: 85% (control), 50%, 25%, and 12.5%. The weight of the trays was checked daily and water was uniformly added to all wells until the target weight was reached.

### Salinity stress

The dry weight and 100% FC saturation of trays was obtained as described above. All trays were maintained at 90% water saturation for a week. The weight of the trays was checked daily, and after seedling establishment (approx. 20 d after germination), water was added containing 0, 50, 100, and 150 mM NaCl. The addition of NaCl solution was uniform to all wells to keep a saturation of 85%.

### Randomization of genotypes for abiotic stress experiment

A total of 16 trays with 14 plants each, were used for this experiment. Genotypes (MIOX4, WT, *vtc*1-1, *vtc*2-1, RV1, RV2) were randomized within the tray to obtain 8 biological replicates for each genotype. These plants were collected and used for RT-qPCR, phenotyping, and photosynthetic measurements. Well number 15 was empty in all trays for water potential meter measurements. Additionally, 8 trays were used for AsA assays; genotypes were randomized for a total of 2 biological replicates. AsA content was determined when the plants were at developmental stage 6.3 for water limitation experiment and at developmental stage 6.1 for salt stress experiment.

### Heat stress

Twelve Quick Pot 15 RW trays were filled with dry growing media Pro-Mix PGX (Promix, PA) (6 trays at 23°C and 6 trays at 37°C) with 14 plants each were used for this experiment. Genotypes (MIOX4, WT, *vtc*1-1, *vtc*2-1, RV1, RV2) were randomized within the tray to obtain 12 biological replicates for each genotype.

Each tray was weighed and recorded. Water was added to each tray until 100% full water capacity. The weights of the trays were checked daily and maintained at 90% water saturation throughout the experiment. Approximately 20 days after germination, the stress was applied.

Plants were treated with a heat shock for two hours at 37 ± 1°C from 11:00 am to 1:00 pm. Control plants were kept at 22 ± 1°C throughout the experiment. Both of the treatments had 65 ± 5% relative humidity and 160-200 µmol m^-2^ s^-1^ light intensity on a long day photoperiod (16 h day: 8 h night). These plants were used for phenotyping, and photosynthetic measurements and leaves were collected for RT-qPCR. Well number 15 was empty in all trays for water potential meter measurements. In addition, 4 trays were used for AsA measurements; genotypes were randomized for a total of 4 biological replicates. AsA content was determined when the plants were at developmental stage 6.1.

### High-throughput phenotyping of Arabidopsis plants

Phenotyping was performed using a Scanalyzer HTS high-throughput phenotyping system using LemnaControl software (Lemnatec, Aachen, Germany) 3 times per week, starting 13-14 days after germination to monitor the vegetative stage through the reproductive stage of plants as described in Acosta-Gamboa *et al.,* 2017. We captured images using visible (RGB), near infrared (NIR), fluorescence (FLUO), and infrared (IR) sensors. Images were analyzed for differences between abiotic stress tolerance treatments and genotypes in the rosette size, leaf shape and area, *in planta* water content, *in planta* chlorophyll fluorescence, and leaf temperature.

### Photosynthetic efficiency measurements

A MultispeQ v1.0 (Kuhlgert *et al*., 2016) was used to measure linear electron flow (LEF), photosynthetic efficiency of photosystem II (ΦII), calculated non-photochemical quenching (NPQ), conductivity of thylakoid membrane to protons (gH+), photon flux (vH+), and electrochromic shift (ECSt). Eight biological replicates for each genotype were measured for salt stress experiment and water limitation stress experiment. For heat stress experiment 12 biological replicates were used.

### Soil water potential meter

Soil water potential (Ψ MPa) was measured with a soil water potential meter (WP4C, Decagon, WA). All measurements were conducted at the same time of day as described (Acosta-Gamboa *et al.,* 2017).

### Real-time quantitative PCR (RT-qPCR) for gene expression analysis

Real-time quantitative PCR (RT-qPCR) was used to quantify expression of genes of interest using reference genes previously published and following MIQE guidelines (Czechowski *et al*., 2005; Udvardi *et al*., 2008). Three biological replicates of leaves were collected at developmental stage 5.0 (Boyes *et al*., 2001). RNA extraction, cDNA preparation, primers efficiency calculation, reference gene validation and RT-qPCR were performed as described (Nepal *et al.,* 2019) with some modifications. Briefly, transcript counts were normalized using internal controls for salt stress (*AtGALDH, UBQ10* and *EF1* α), heat stress (*EF1*α and *AtGALDH*), and water limitation (*Actin 2, AtGALDH* and *EF1* α) (Supplementary Table 1). The relative expression of the genes was calculated using the Biogazelleqbase+ software (Version 3.2). Three biological replicates and two technical replicates were used for RT-qPCR.

### Seed yield

The weight of 100 seeds per treatment was determined 3 times for each genotype for each treatment, and the average was used to determine the total number of seeds per treatment.

### Seed germination

Germination rates were scored as germinated seedlings versus total seeds by placing 25 seeds on MS media supplemented with kanamycin for 11-15 days.

### Statistical analyses

For statistical evaluation of the experiments, the GRAPH PAD Prism 8 (Version 8.0.1, 2019) was used. Variation in photosynthetic efficiency parameters, AsA measurements, water potential meter readings, seed germination, seed yield, and RT-qPCR were analyzed using ANOVA comparing the mean against the control. To correct for multiple comparisons, Dunnett’s test was used at α=0.05 Data presented are means ± standard error. The variation in color classification was analyzed using ANOVA in R (Version .3.5.2, 2018). To correct for multiple comparisons, Tukey’s post-hoc test was used at α=0.05. To analyze projected leaf surface area and compactness, we performed an analysis of repeated measures for split plot design with the days of 21, 23, 25, and 28 when the stress factors were applied. To build a linear mixed effect model, trays and plants were treated as random effects, and the variance and covariance across the days were incorporated into parameter estimation. Tukey’s multiple comparison testing among six genotypes was performed at 0.05 level of significance.

## Results

The MIOX4 over-expresser line selected for this study (L21) is a homozygous line and contains a single copy of the MIOX4 ORF. This line has high AsA content, possesses tolerance to various abiotic stresses, and displays increased biomass (Lisko *et al*., 2013; Yactayo-Chang, 2011; Yactayo-Chang *et al*., 2018; Nepal *et al*., 2019). The Lorence laboratory has used this line for over 7 years without gene silencing issues.

The selection of the *vtc* mutants was to account for the contribution of two pathways present in the metabolic network of AsA production. The *vtc*1-1 gene encodes the enzyme involved in catalyzing the conversion of d-mannose-1-P to GDP-mannose, a step not only present in the d-mannose/l-galactose pathway, but also in the l-gulose pathway (Wolucka and Van Montagu 2003). In *vtc*2-1, there is an encoded mutation in the GDP-L-galactose phosphorylase, which results in complete loss of its activity and accumulates only 20-30% of the WT AsA levels (Conklin *et al*., 2000). This mutant is only affected in the d-mannose/l-galactose pathway.

### MIOX4 gene expression and ascorbate are elevated under water limitation stress

Under severe water limitation, the normalized expression of the *MIOX4* gene was significantly higher in RV1, RV2, and MIOX4 when compared with WT. This expression was also relatively higher when the available water was reduced (12.5%) (Figure 4A). Under normal water saturation, ascorbic acid content was significantly higher in MIOX4 and the restored lines, and significantly lower in the *vtc* mutants compared to WT (Figure 4B). When water availability was reduced to 12.5%, ascorbate content in MIOX4 and RV2 were significantly higher compared to WT (Figure 4C). In both normal and stress conditions, the *vtc* mutants held the lowest ascorbate content. These results indicate that MIOX4 increased the ascorbate content in restored lines under normal and water limitation conditions.

**Figure 1.**
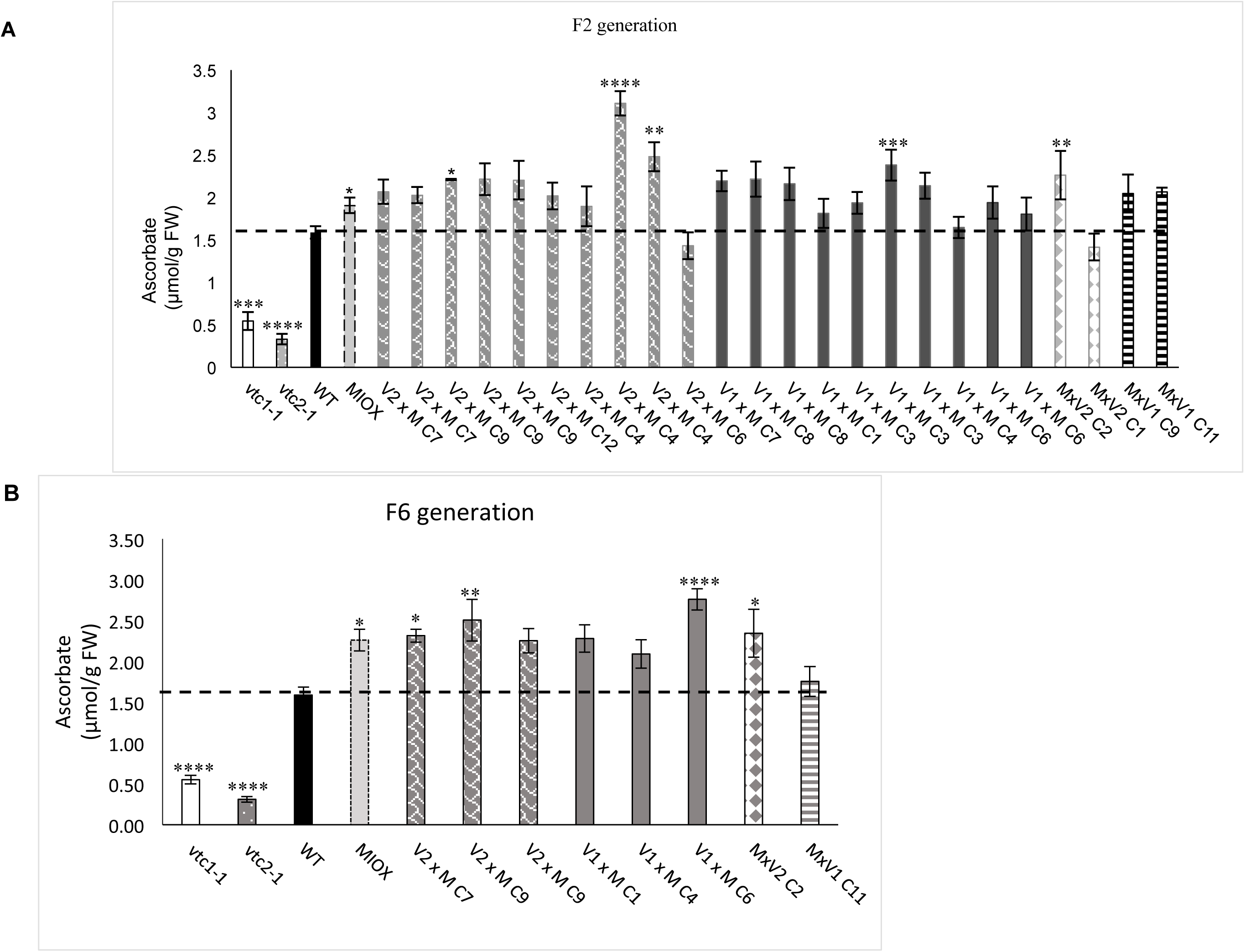
Total foliar ascorbate content of MIOX4 x *vtc* crosses and control lines. **(A)** Ascorbate content of F2 generation crosses. **(B)** Ascorbate content of F6 generation homozygous crosses. “C” corresponds to the cross number. Data are means ± SEM (n=5). Asterisks represent significant differences (p < 0.0001).

**Figure 2.**
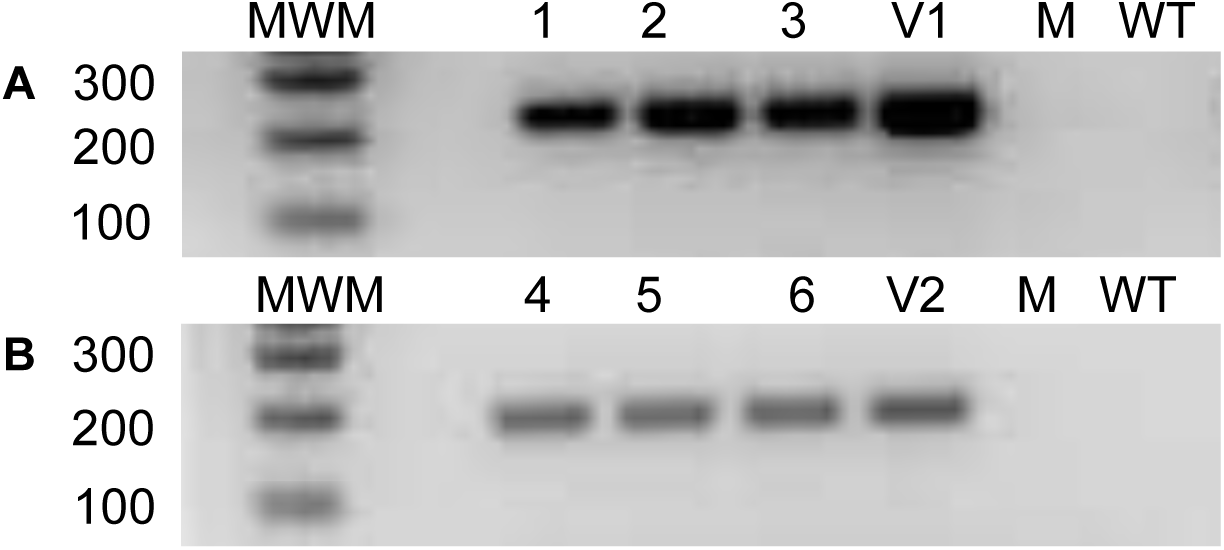
(A) RT-PCR and **(B)** PCR analysis for the MIOX4 x vtc crosses and controls. M: *MIOX*4, WT: wild type, V1: *vtc*1-1 and V2: *vtc*2-1, MWM =molecular weight marker. Samples 1-6 are as described in Table 1.

**Figure 3.**
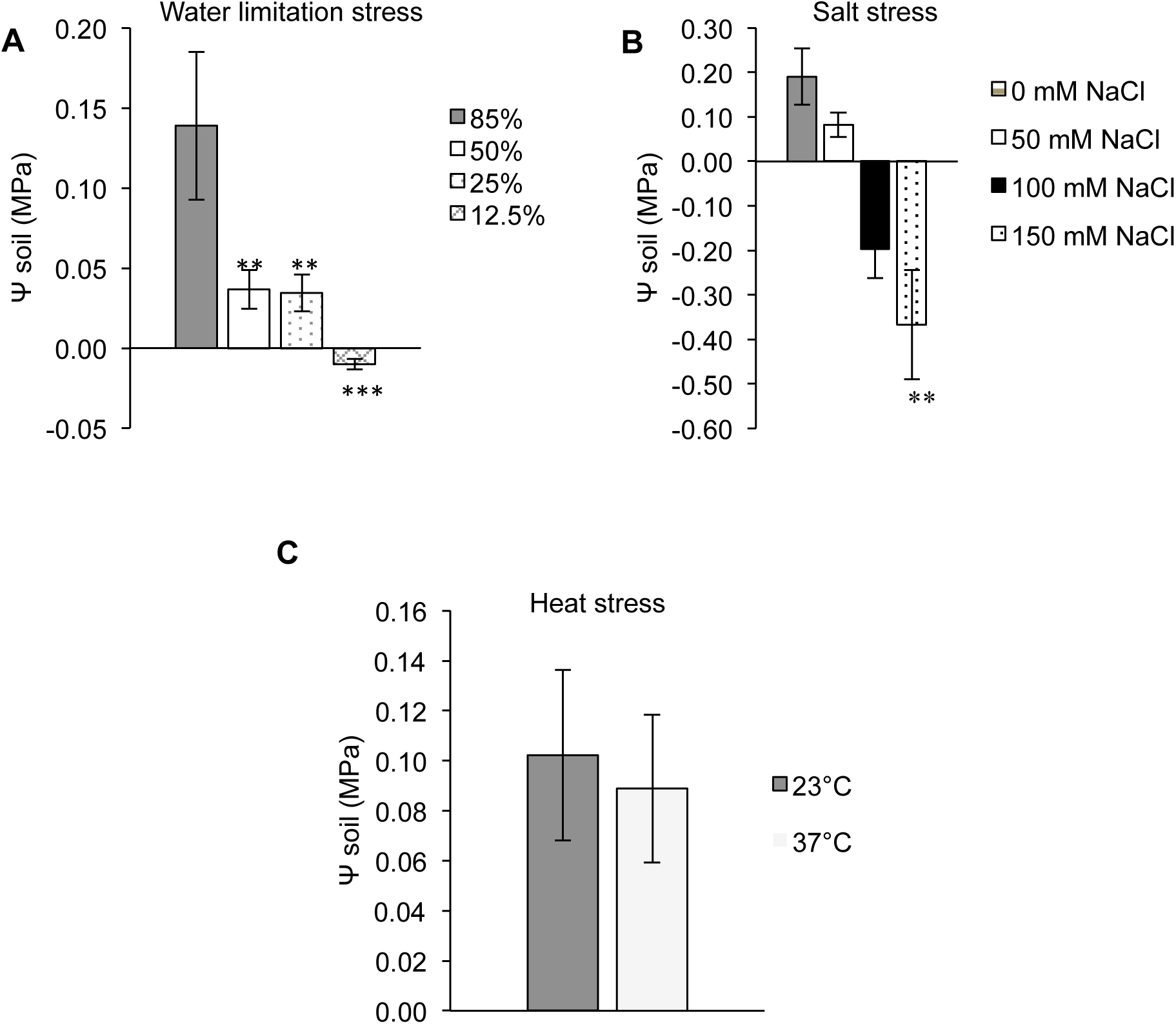
Soil water potential measurements. **(A)** Effect of water deficit on soil water potential. **(B)** Effect of salt application on soil water potential. **(C)** Effect of heat stress on soil water potential. Data are means ± SEM (n =5). Asterisks represent significant differences (p < 0.0001).

**Figure 4.**
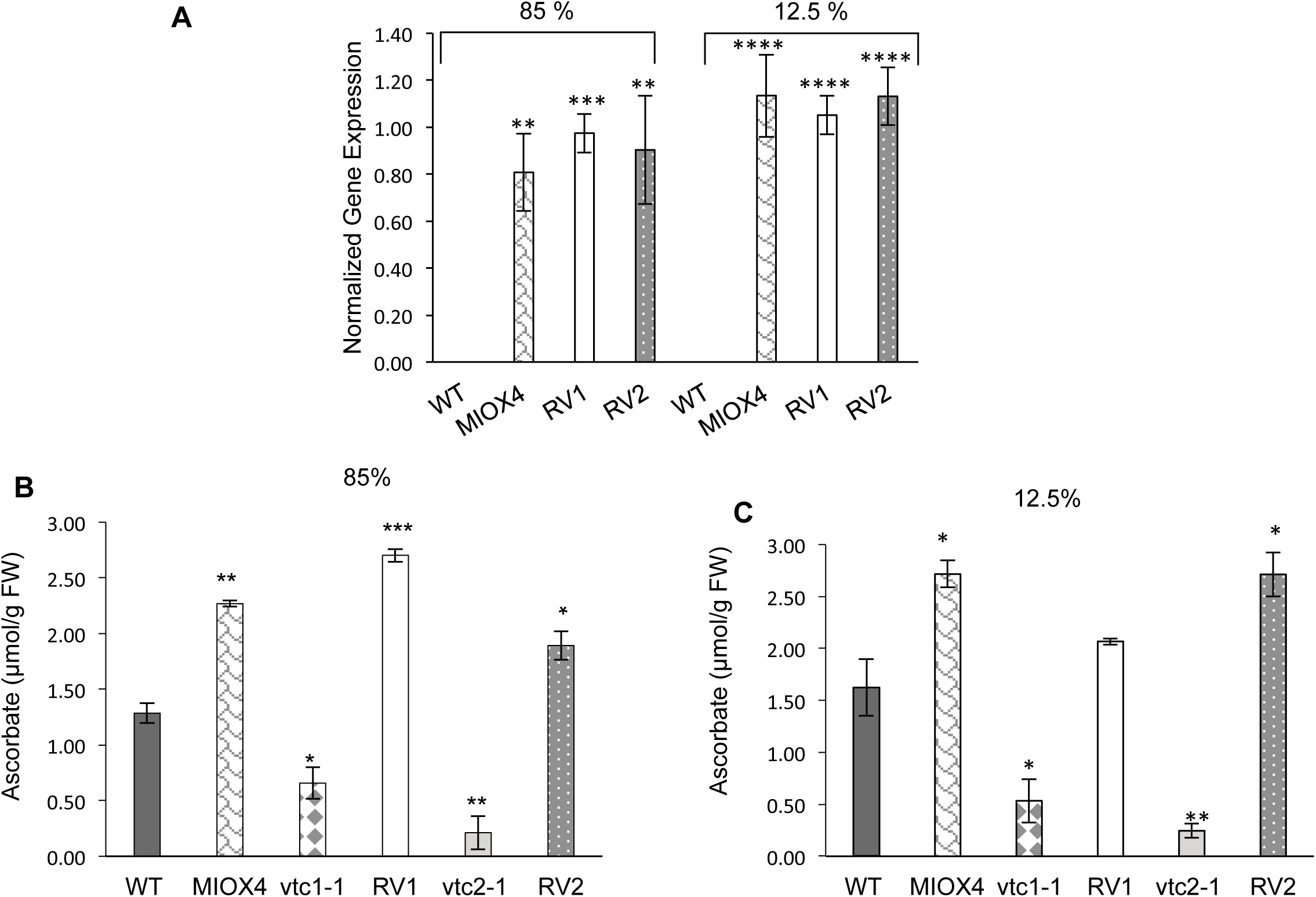
MIOX4 gene expression and foliar ascorbate content of the MIOX4 x *vtc* crosses in response to water limitation stress. **(A)** Expression of *MIOX4* quantified via RT-qPCR. Expression was normalized to *Actin* 2, *EF1*α, and *AtGALDH.* Ascorbate content of MIOX4 x *vtc* crosses under normal (**B**) versus severe water limitation (**C**). Data are means ± SEM (n=3). Asterisks represent significant differences *p<0.05, ** p<0.01, and ***p<0.001.

### MIOX4 restores the biomass in *vtc* mutants under water limitation stress

Water limitation negatively affects the plant growth and development. Interestingly biomass was clearly affected under severe water limitation (12.5% and 85% FC). When analyzing the RGB images, signs of chlorosis or necrosis were observed under water limitation and were absent in control (Figure 5A-B). The growth of plants was significantly affected by the water limitation stress (Supplementary Table 2A). When plants were grown under normal water availability, the restored lines grew better compared with their respective controls (*vtc*1-1 and *vtc*2-1) and bigger than MIOX4 or WT (F_5,42_=12.708, p<0.001) (Figure 5C, Supplemental Table 2D). Twenty-eight days after germination, at the end of phenotyping, *vtc* mutants recorded the lowest projected leaf surface area, compared with the rest of the genotypes (Figure under 85% water saturation (Figure 5C), and under 12.5% water limitation (F_5,42_=12.291, p<0.001) (Figure 5D, Supplementary Table 2C). Plants under severe water limitation were more compact than plants grown under normal conditions, and at the end of the experiment, the leaves started to curl down (compactness measurements) (Figure 5E-F). These results indicate that plants limited their growth rate and biomass under water limitation stress. Restored lines had enhanced biomass compared to parent lines and WT control under water limitation stress, suggesting that *MIOX4* restored the biomass of restored lines under normal and water limitation stress.

**Figure 5.**
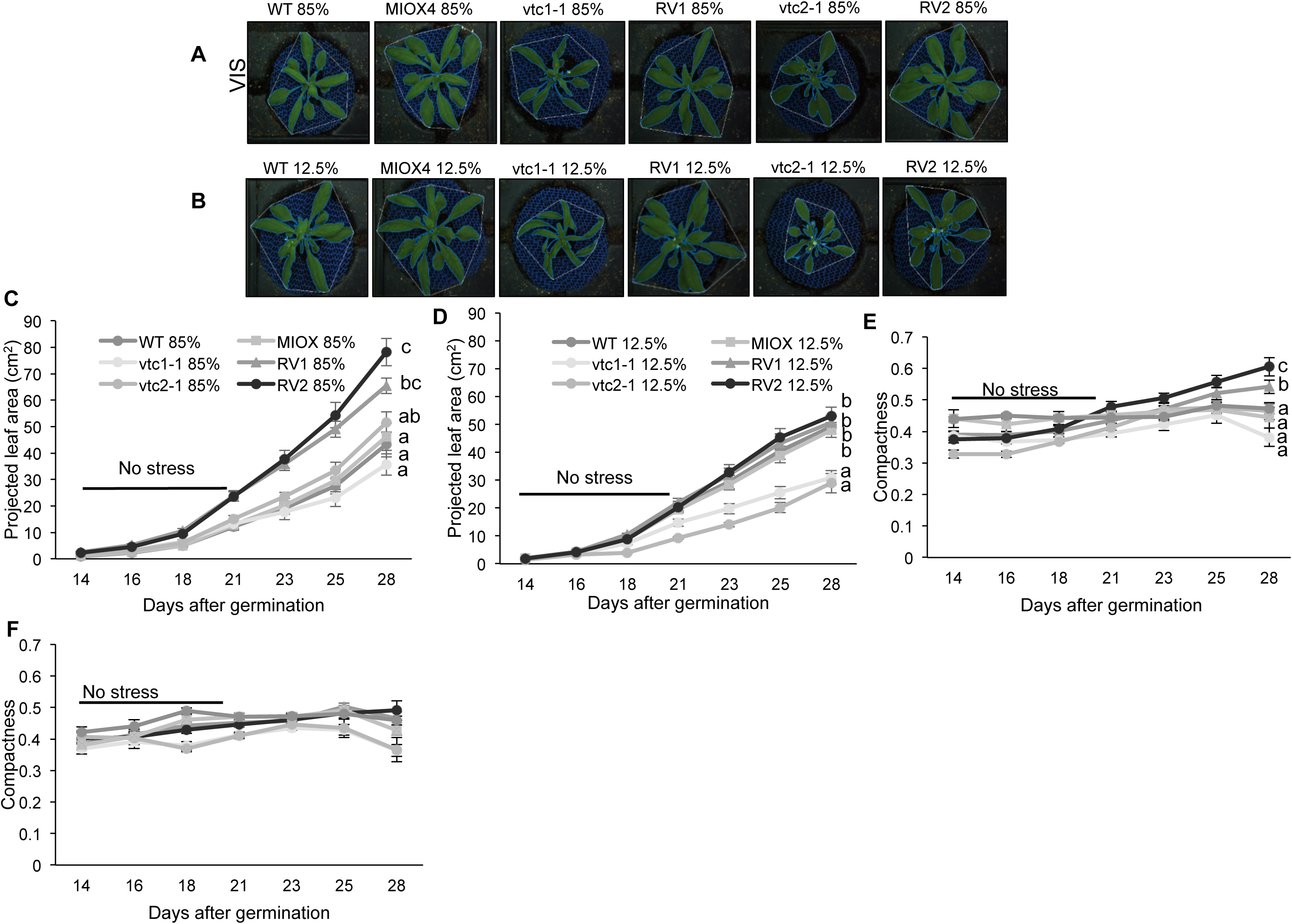
Phenotype and growth of MIOX4 x vtc crosses and controls under water limitation conditions. Representative images of plants growing under control (**A**), versus severe water limitation (**B**) conditions. Total projected leaf area of plants under control (**C**) versus 12.5% water limitation (**D**). Compactness [total leaf area/convex hull area] of plants under control (**E**) versus severe water limitation (**F**). Data are means ± SEM (n =8). Different letters represent significant differences between treatments (p < 0.0001).

### *In-planta* relative water content is lower in *vtc* mutants under water limitation stress

Relative water content, relative chlorophyll content and relative chlorophyll fluorescence were obtained from NIR, VIS and FLUO sensors respectively and impacted by water limitation stress in time dependent manner (Supplementary Figure 2 and 3). Under normal conditions chlorosis was higher in *vtc*-mutants compared to MIOX4, WT and restored lines at 28 days after germination, last days of phenotyping (Figure 6A). Under 12.5% water limitation stress, chlorosis was higher in *vtc2-1* mutant and RV2 lines compared to other genotypes (Figure 6D). Similarly, *in-planta* relative water content was lower in *vtc* mutants compared to other genotypes under normal condition (Figure 6B), and under 12.5% water limitation stress (Figure 6E, Supplementary Figure 4A, 4C) at 28 days after germination. Furthermore, under normal conditions, the relative chlorophyll fluorescence was similar in all genotypes (Figure 6C, Supplementary Figure 4B). Interestingly, relative chlorophyll fluorescence was higher in *vtc* mutants and WT compared to MIOX4 and restored lines under 12.5% water limitation stress (Figure 6F, Supplementary Figure 4D). These results indicate that, the relative chlorophyll content was restored in restored lines under normal condition, and *in-planta* relative water content was restored in restored lines under normal conditions and water limitation stress condition. The *vtc* mutants and WT lines were more stressed compared to restored lines and MIOX4 line under water limitation stress.

**Figure 6.**
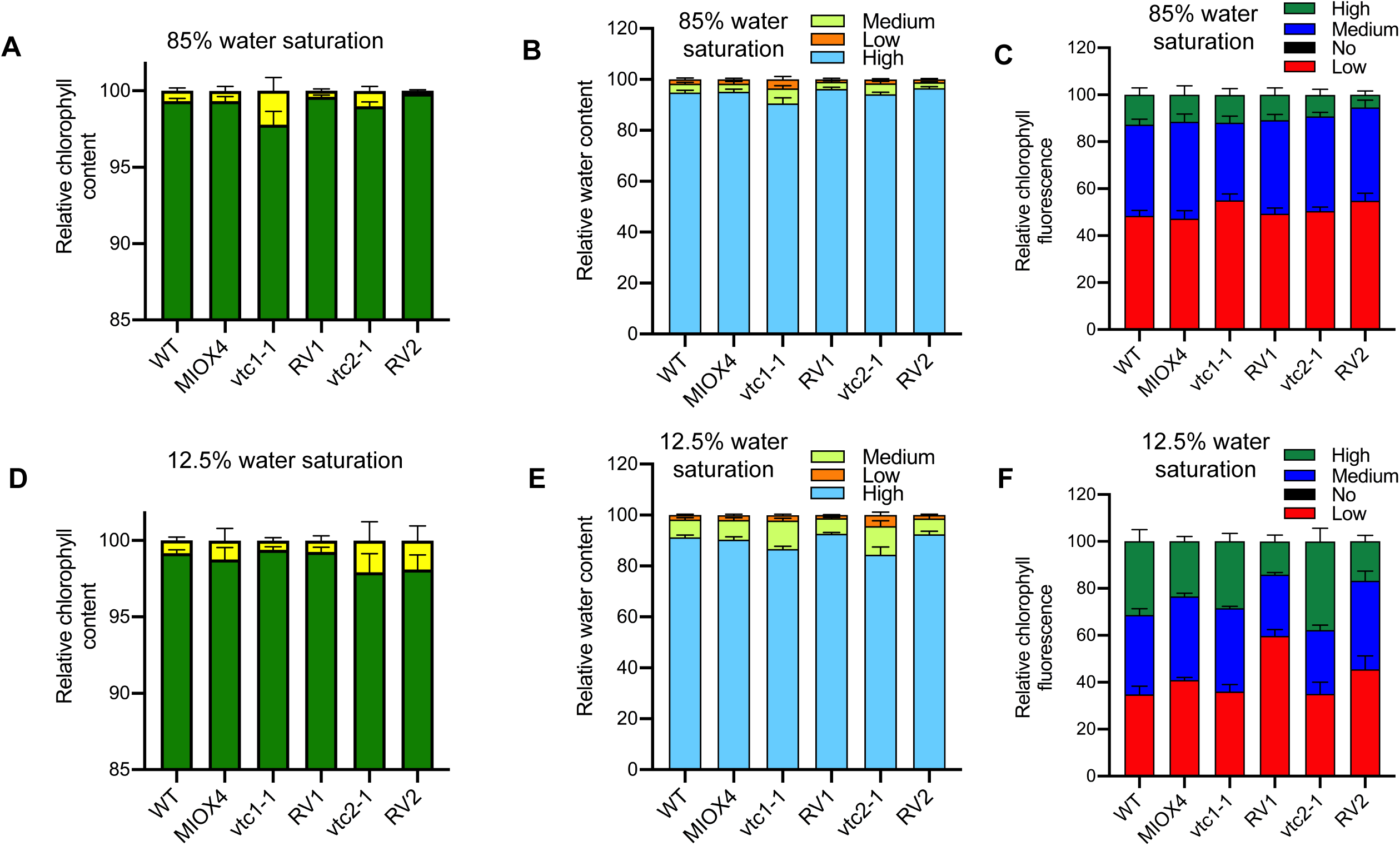
Chlorosis, *in planta* water content and *in planta* chlorophyll fluorescence of MIOX4 x vtc crosses in response to water limitation stress. Chlorosis of plants grown under control (**A**) versus severe water limitation conditions (**D**). *In-planta* relative water content of plants grown under control (**B**) versus 12.5% water saturation (**E**). *In planta* chlorophyll fluorescence of plants grown under control (**C**) versus12.5% water saturation (**F**). Observations were recorded 28 days after germination. Data are means ± SEM (n=8).

### Seed yield decreased under water limitation stress

The yield and yield components were affected by the abiotic stresses. Under water limitation, seed yield decreased 1.5 to 2 fold when compared with 12.5% and 85% in all genotypes (Figure 7).

**Figure 7.**
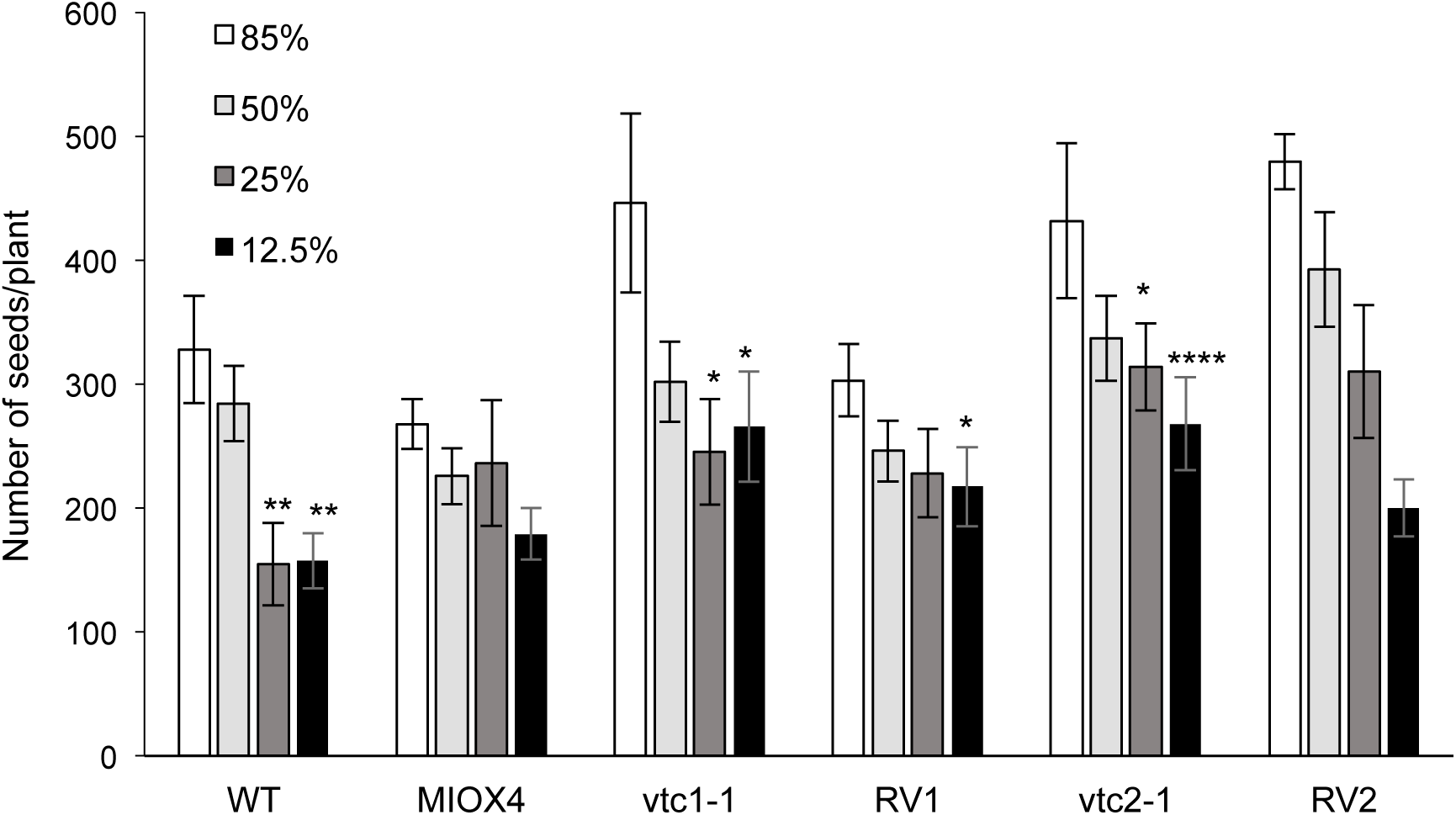
Seed yield of MIOX4 x vtc crosses in response to water limitation stress. Data are means ± SEM (n=8). Asterisks represent significant differences *p<0.05, **p<0.01, and ****p<0.001.

### *MIOX4* gene expression and ascorbate is elevated under salt stress

Under salt stress conditions, normal condition (0 mM NaCl) and 150 mM NaCl application, expression of *MIOX4* was significantly higher in MIOX4 and restored lines compared to WT (Figure 9A). Under 0 mM NaCl, AsA content was significantly lower in *vtc* mutants, and significantly higher in MIOX4 and restored line compared to WT (Figure 8B). Similarly, under 150 mM NaCl treatment, ascorbate content was significantly higher in MIOX4 and restored lines, and significantly lower in *vtc2-1* mutant compared to WT (Figure 8C). In both normal and salt stress condition, the *vtc* mutant lines had lowest AsA content and restored lines had high ascorbate. These results indicate that MIOX4 restored the AsA content in restored lines under normal and salt stress conditions.

**Figure 8.**
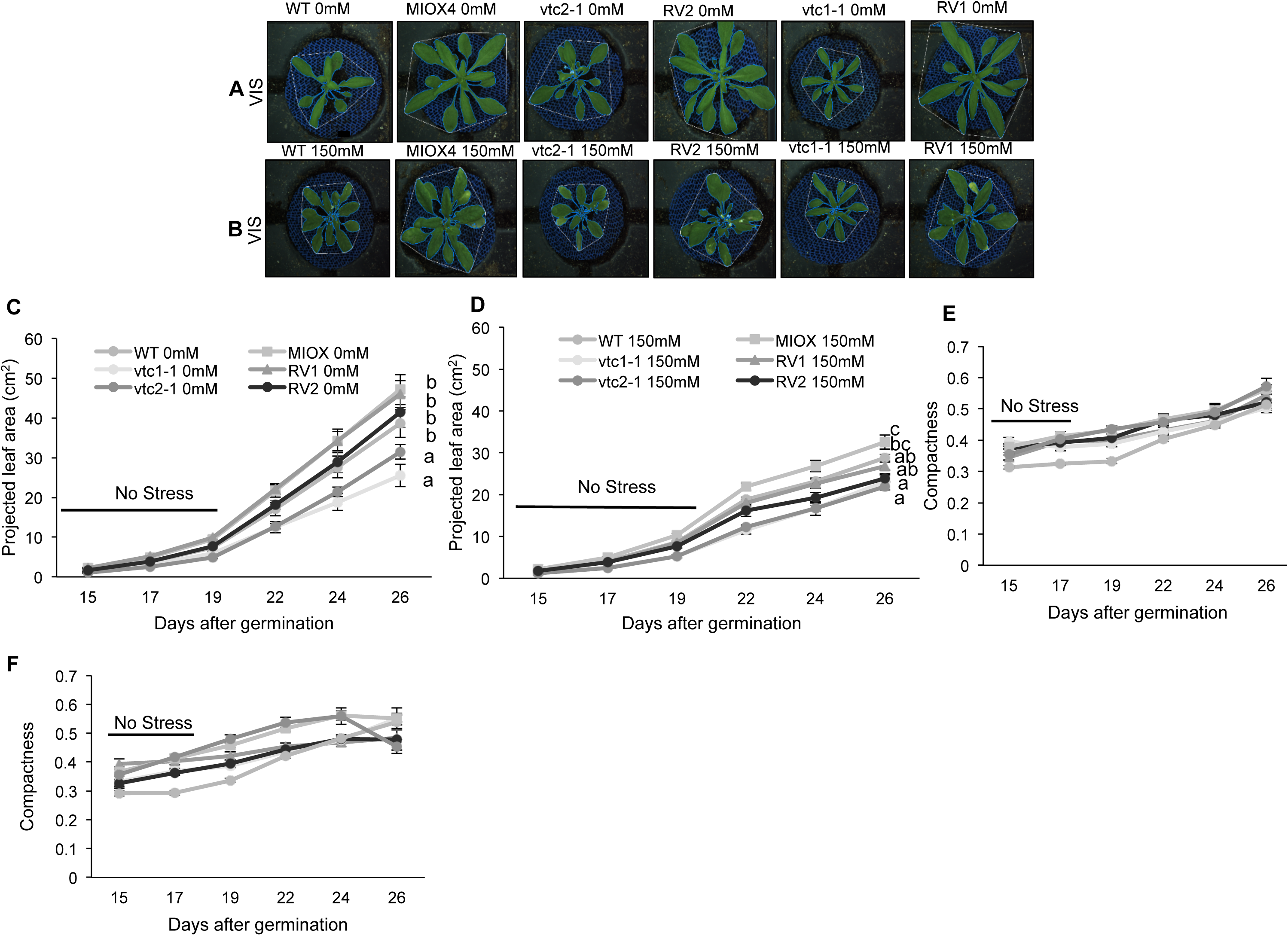
Phenotype and growth of MIOX4 x vtc crosses and controls under salt stress. Representative images of plants growing under control (**A**), 150 mM NaCl (**B**) conditions. Total projected leaf area of plants under control (**C**) versus 150 mM NaCl (**D**). Compactness [total leaf area/convex hull area] of plants under control (**E**) versus severe water limitation (**F**). Data are means ± SEM (n =8). Different letters represent significant differences between treatments (p < 0.0001).

**Figure 9.**
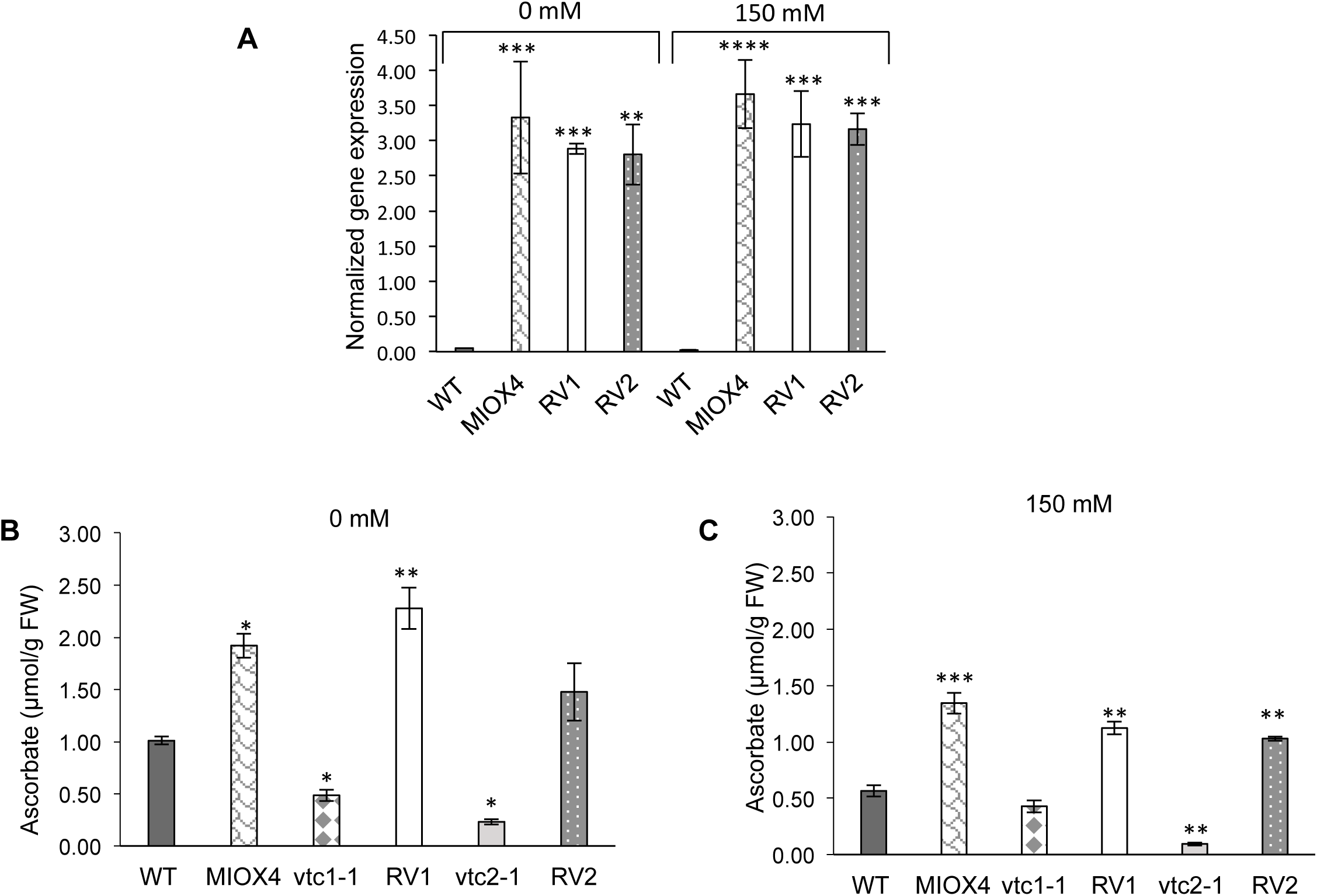
MIOX4 gene expression and foliar ascorbate content of the MIOX4 x *vtc* crosses in response to salt stress. **(A)** Expression of *MIOX4* quantified via RT-qPCR. Expression was normalized to *Actin* 2, *EF1*α, and *AtGALDH.* Ascorbate content of MIOX4 x vtc crosses under normal (**B**) versus salt stress (**C**). Data are means ± SEM (n=3). Asterisks represent significant differences * p<0.05, ** p<0.01, and *** p<0.001.

### Restored lines have enhanced biomass compared to *vtc* mutants

Salt application caused a penalty in plant growth and development over time. Biomass was affected under the salt stress regimes (0 mM and 150 mM NaCl) (Supplementary Table 2B). When analyzing the RGB images, signs of chlorosis or necrosis were observed when 150 mM NaCl stress and were absent in control conditions (Figure 9A-B). Under the highest application of sodium chloride, the MIOX4 line was statistically bigger than the restored and mutant lines (F_5,42_=9.759, p<0.001) (Figure 9D, Supplementary Table 2F). The mutant lines (*vtc*1-1 and *vtc*2-1) were statistically smaller under no stress, compared with the rest of the genotypes (F_5,42_=6.687, p<0.001) (Figure 9C, Supplementary Table 2E). Compactness showed that plants under stress at the end of their life cycle tend to curl their leaves, compared with no salt conditions (Figure 9E-F).

### Chlorosis is lower in *vtc* mutants under salt stress condition

Salinity stress reduces chlorophyll content in plants. Under 0 mM NaCl, the chlorophyll content was similar in all genotypes (Figure 10A). Under 150 mM NaCl, the chlorosis was higher in all genotypes compared to the plants grown in 0 mM NaCl. Similarly, the chlorosis was lower in *vtc* mutants compared to restored lines, MIOX4 and WT (Figure 10D). The *in planta* relative water content was similar in all genotypes under 0 mM NaCl treatment (Figure 10B, Supplementary Figure 5A, 5C). Under 150 mM NaCl treatment, *in-planta* relative water content was lower in *vtc* mutants compared to restored lines, MIOX4 and WT (Figure 10E, Supplementary Figure 5C). The relative chlorophyll fluorescence was similar in all genotypes under 0 mM and 150 mM NaCl treatments (Figure 10C-F, Supplementary Figure 5B, 5D).

**Figure 10.**
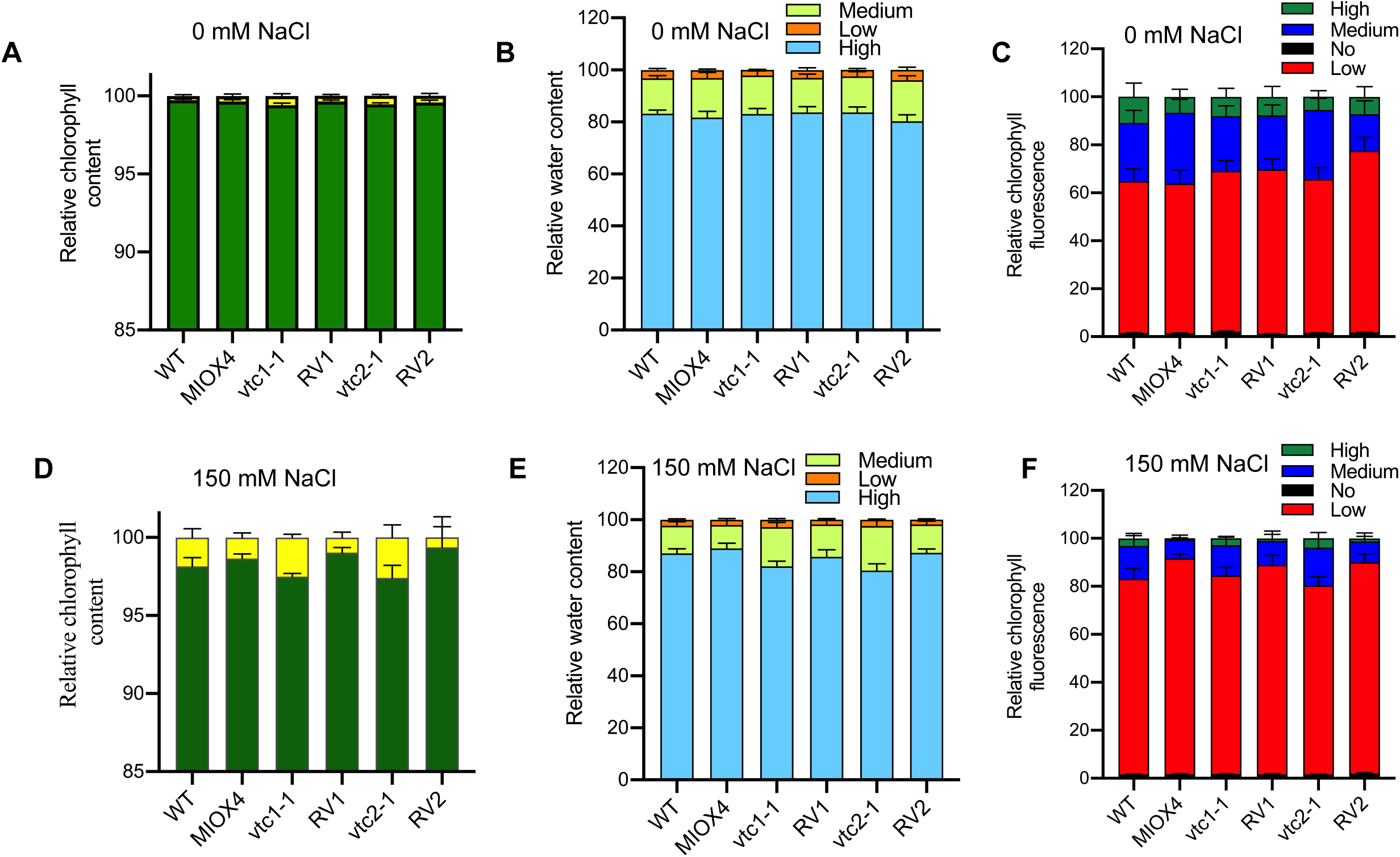
**Chl**orosis, *in planta* water content and *in planta* chlorophyll fluorescence of MIOX4 x *vtc* crosses in response to salt stress. Chlorosis of plants grown under control (**A**) versus 150 mM NaCl (**D**). *In-planta* relative water content of plants grown under control (**B**) versus 150 mM NaCl (**E**). *In planta* chlorophyll fluorescence of plants grown under control (**C**) versus 150 mM NaCl (**F**). Observations were recorded 28 days after germination. Data are means ± SEM (n=8).

### Seed yield under salt stress condition

In the case of salinity, plants treated with 50 mM tended to produce a higher number of seeds per plant compared with the rest of the treatments. However, when salt application increased (up to 150 mM NaCl), seed yield was dramatically decreased (Figure 11).

**Figure 11.**
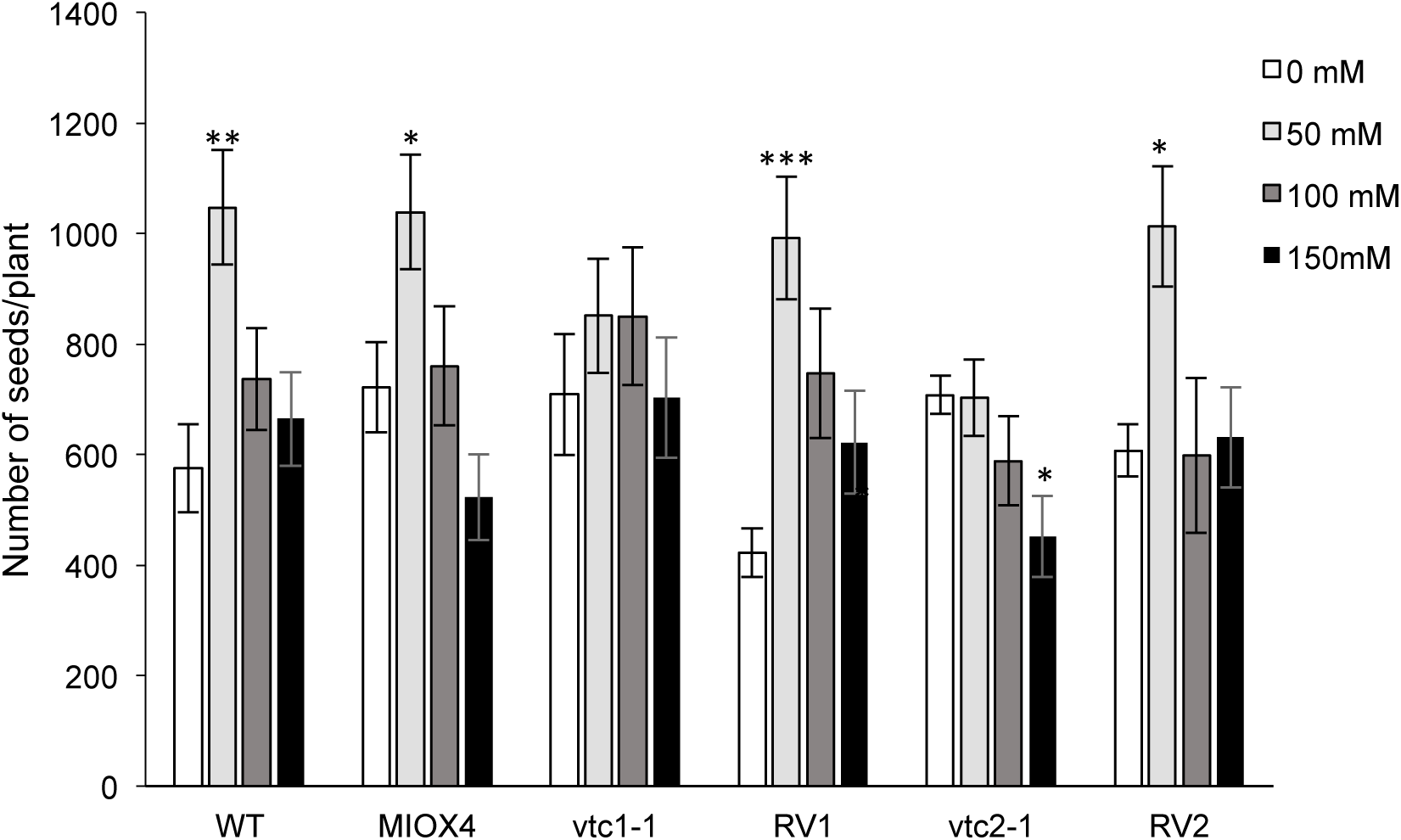
Seed yield of MIOX4 x *vtc* crosses in response to salt stress. Data are means ± SEM (n=8). Asterisks represent significant differences; * p<0.05, ** p<0.01, and *** p<0.005.

### *MIOX4* gene expression and ascorbate is elevated under heat stress

Under the heat shock, at 37°C *MIOX4* gene expression was higher in MIOX4 and restored lines compared to the plants grown at 23°C. Furthermore, at 37°C or 23°C *MIOX4* expression was higher in MIOX4 and restored lines compared to WT (Figure 12A). The *vtc* mutant lines presented the lowest AsA content under normal and stressed conditions. Under no stress, RV1 and MIOX4 produced the highest ascorbate. When the heat shock was applied, the three lines over-expressing the MIOX4 ORF displayed the highest ascorbate content (Figure 12B-C).

**Figure 12.**
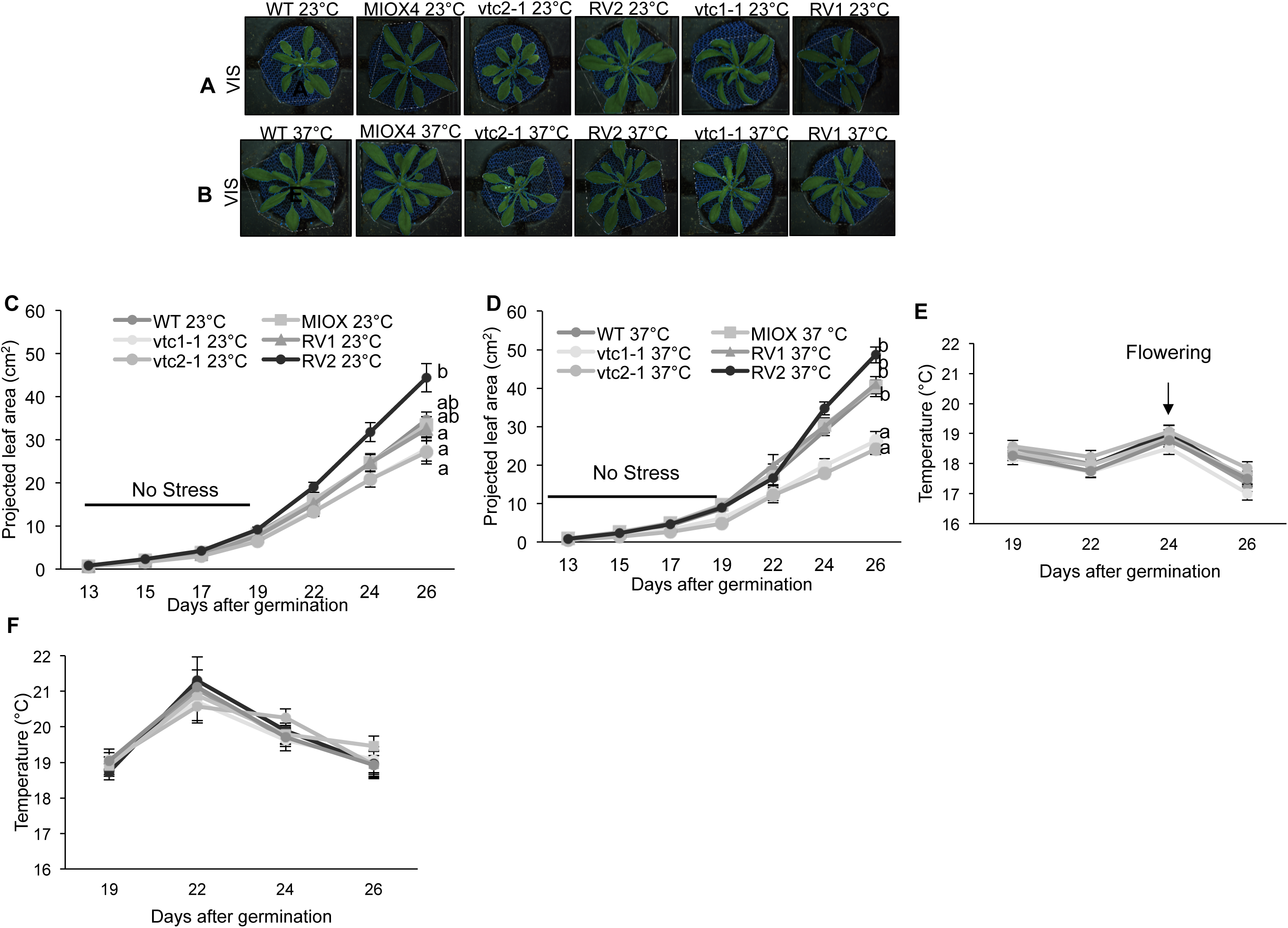
Phenotype and growth of MIOX4 x *vtc* crosses and controls in response to heat stress. Representative images of plants growing under control (**A**), versus heat (**B**). Total projected leaf area of plants under control (**C**) versus heat (**D**). Leaf temperature of plants grown under control (**E**) versus heat conditions (**F**). Data are means ± SEM (n =8). Different letters represent significant differences between treatments (p < 0.0001).

### Heat stress affects the rosette growth at advance stages of plant development

An increase in fourteen-Celsius degrees for two hours affected the performance of the genotypes. The mutant lines grew the poorest, and the RV2 line displayed the highest projected leaf area under normal (continuous 23°C) and heat stress (Figures 13A-D). The IR sensor was able to detect an increase in temperature when the plants began flowering (normal conditions) (Figure 13E). One day after the heat shock was applied (22 days after germination), the sensor detected an increase in leaf temperature compared with all other days (Figure 13F, Supplementary Figure 6C, 6F).

**Figure 13.**
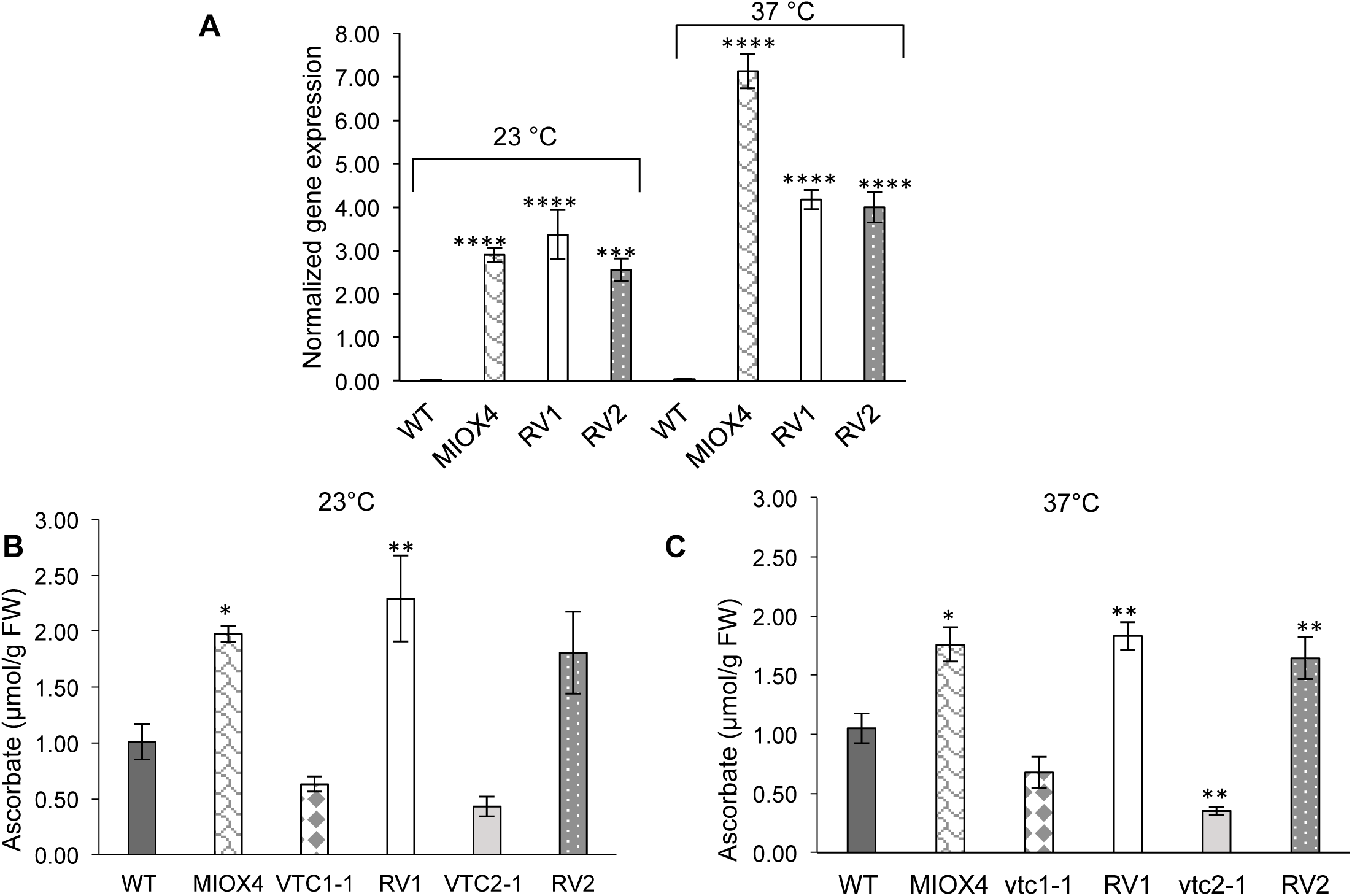
MIOX4 expression and foliar ascorbate content of the MIOX4 x *vtc* crosses in response to heat stress. **(A)** Expression of *MIOX4* quantified via RT-qPCR. Expression was normalized to *Actin* 2, *EF1*α, and *AtGALDH.* Ascorbate content of MIOX4 x *vtc* crosses under normal (**B**) versus heat stress (**C**). Data are means ± SD (n=3). Asterisks represent significant differences * p<0.05, ** p<0.01, and *** p<0.001.

### Photosynthetic efficiency was slightly affected by the stresses

The stresses applied have shown negative affect on photosynthetic efficiency (PE), chlorophyll fluorescence, and linear electron flow (LEF). The photosynthetic parameters (ΦII, NPQt, vH+, gH+, ECSt; not shown) were not significantly different in plants grown under water limitation and heat stresses. However, most of them showed a trend in which, LEF was higher and NPQt was lower in normal conditions (85% and 0 mM NaCl), compared with the stress treatments (12.5% and 150 mM NaCl). Under heat stress, at 37°C *vtc*1-1 showed the lowest LEF and highest NPQt at 24 days after germination (Figures 14A, 14C) compared to other genotypes. At 23°C, LEF was significantly higher and NPQt was significantly lower in RV2 compared to the rest of the Arabidopsis lines at 24 days after germination (Figures 14B, 14D).

**Figure 14.** Effect of heat treatment on *A. thaliana* photosynthetic efficiency. **(A)** and **(C)** Linear electron flow as an indicator of photosynthetic efficiency during heat stress conditions. **(D)** and **(E)** Non-photochemical quenching during heat stress conditions. Data were obtained 24 days after germination. Data represents the means ± SEM (n=8).

### Relative chlorophyll content, *in-plant* relative water content, and relative chlorophyll fluorescence under heat stress

Under the heat stress condition, there was no significant difference in relative chlorophyll content was observed in all genotypes (Figures 15A, 15D). Similarly, no significant difference was observed among genotypes in *in-planta* relative water content under heat stress conditions (Figure 15B, 15E, Supplementary Figure 6B, 6D). Furthermore, there was no significant difference observed among genotypes in relative chlorophyll fluorescence under heat stress condition (Figure 15C, 15F, Supplementary Figure 6A, 6C).

**Figure 15.**
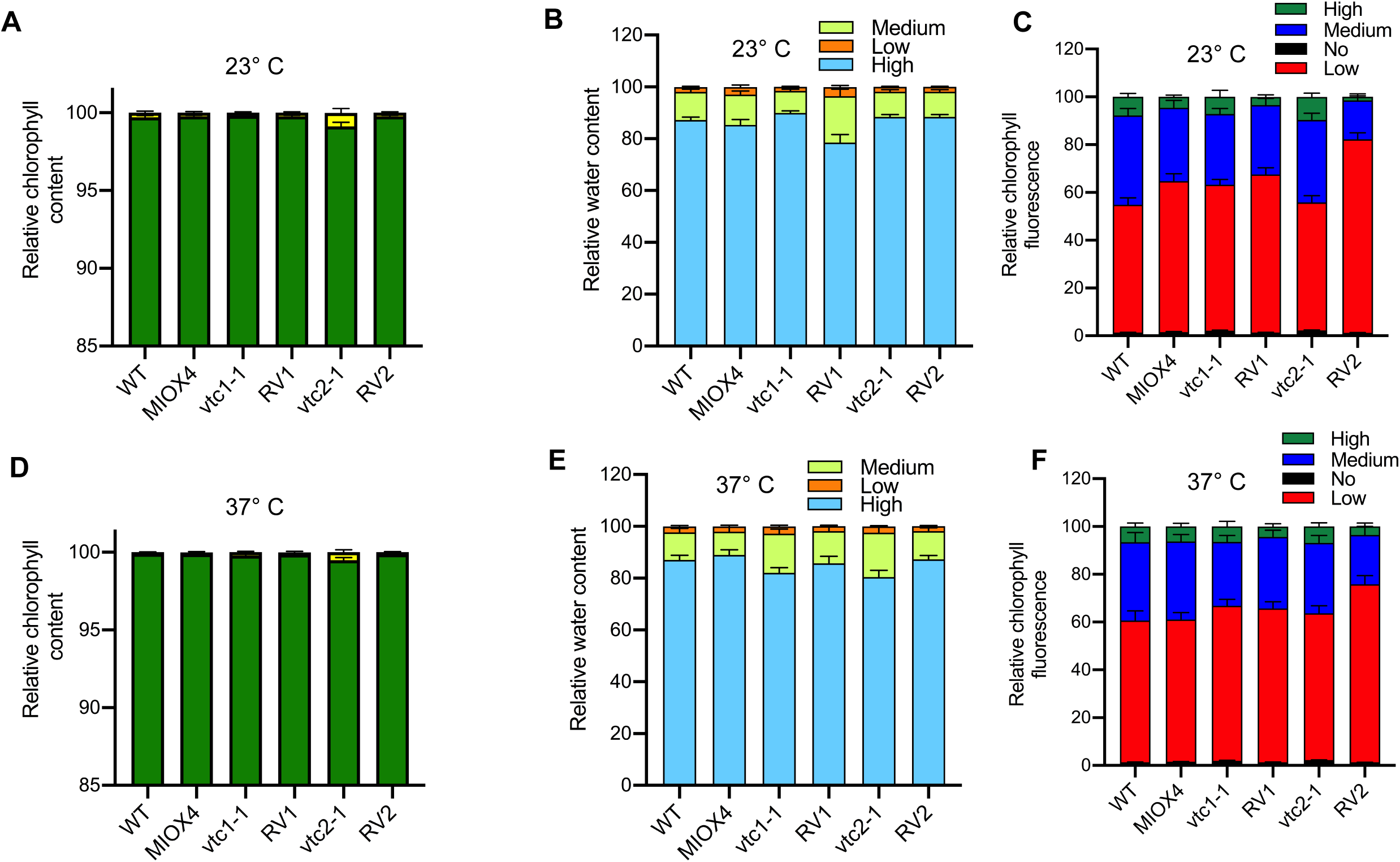
Chlorosis, *in planta* water content and *in planta* chlorophyll fluorescence of MIOX4 x *vtc* crosses in response to heat stress. Chlorosis of plants grown under control (**A**) versus heat (**D**). *In-planta* relative water content of plants grown under control (**B**) versus heat (**E**). *In planta* chlorophyll fluorescence of plants grown under control (**C**) versus heat (**F**). Observations were recorded 28 days after germination. Data are means ± SEM (n=8).

## Discussion

Ascorbate content has long been studied in order to determine its significance as an antioxidant and in plant growth and development. In order to understand which pathway drives ascorbate production, some scientists have used feeding assays (Davey *et al*., 1999; Barata-Soares *et al*., 2004; Creameans *et al*., 2017) or biochemical and molecular enzyme identification/characterization techniques (Imai *et al*., 1998; Conklin *et al*., 1999*a*; Gatzek *et al*., 2002; Laing *et al*., 2007). Here, we have explored the increased production of AsA by constitutive over-expression of one enzyme present in the *myo*-inositol pathway and the restoration of production of this small molecule using low vitamin C lines as background.

### MIOX4 over-expression restores AsA production via the MI pathway

A recent study has disputed the MI pathway’s contribution to AsA production (Kavkova *et al*., 2018), but we have shown that under normal and abiotic stresses conditions, this route contributes to the AsA production in Arabidopsis restored lines. The published evidences indicate that MIOX4 overexpression increases AsA and tolerance to abiotic stresses (Lorence *et al*., 2004; Donahue *et al*., 2010; Tóth *et al*., 2011; Alford *et al*., 2012; Duan *et al*., 2012; Lisko *et al*., 2013: Nepal *et al.,* 2019; Acosta-Gamboa *et al.,* 2020). Our measurements show that, indeed, by over-expressing this specific gene, plants produced more AsA compared with controls. This phenomenon was also observed in the *vtc* mutant lines crossed with the MIOX4 line under normal conditions (Figures 1A-B) and under abiotic stress (Figures 4C, 8C, 12C). When working with transgenic plants with the 35S constitutive promoter, the chances of silencing are increased if the Arabidopsis line is not homozygous (Mlotshwa *et al*., 2010). In this study, we rated seed germination in media with selectable marker kanamycin to obtain homozygous restored lines (Supplementary Figure 1).

Our results show that MIOX4 restored the function of the MI pathway and increased the production of AsA. RV1 and RV2 lines displayed 5 and 8 fold increase in AsA content compared with controls (Figure 1B). Ascorbate content in the *vtc* mutants was comparable to the levels detailed by other scientists (Conklin *et al*., 1996, 1997, 2000; Müller-Moulé *et al*., 2004; Dowdle *et al*., 2007; Lim *et al*., 2016). The presence of the *vtc*1-1 and *vtc*2-1 single point mutation in the restored lines supported this result (Figure 2, Table I). For these two mutants, the contribution of AsA from the Smirnoff/Wheeler and l-gulose pathway is knocked down (Conklin *et al*., 2000). In agreement with this, our data shows that the MI pathway plays a crucial role in restoring AsA production when the Smirnoff/Wheeler and l-gulose pathways are malfunctioning.

Plants subjected to abiotic stresses such as high light, water limitation, and low/high temperatures produce more ROS (Foyer and Noctor, 2009). Under these conditions, ascorbate acts as a central component in regulating ROS levels (Foyer and Noctor, 2011). In order to determine if the increase in AsA made a significant impact on abiotic stress tolerance, we applied three different abiotic stresses to the *A. thaliana* lines (WT, MIOX4 over-expresser, restored lines and *vtc* mutants).

### Impacts of abiotic stresses on plant growth and development: understanding the physiology with high-throughput phenotyping

Applying abiotic stresses, such as water limitation or salt, requires precise and well-monitored experimental design; this includes controlling the soil water content and defining the time to start the stress, as plant response is dependent on the developmental stage (Skirycz *et al*., 2010; Verelst *et al*., 2010). Soil water potential measurements can also standardize the quantification of water added to each of the treatments and contribute to experimental reproducibility. Our measurements show water potential values that are consistent with moderate to severe soil water deficit (Durand *et al*., 2016). When the salt was applied, soil water availability decreased due to a difference in osmotic potential of the soil matrix (Lamsal *et al*., 1999). In agreement with these observations, our data shows a reduction in water availability when salt increased, and under water limitation (Figure 3A-B).

When water availability is reduced due to salt application, water limitation, or heat, plants are negatively affected. Processes such as membrane disruption, metabolic toxicity, reduction in photosynthetic efficiency, and increased ROS production occur and can lead to plant death (Yeo, 1998; Mcdowell *et al*., 2008; Hasanuzzaman *et al*., 2013). Our data suggests that the negative effects caused by each of the stresses were time and treatment dependent (Figures 5, 9, 13). Plant plasticity is complex, and the use of different sensors helps understand the physiology of the plant. A variety of sensors are available to capture signals from the visible, fluorescent, and infrared light spectra (Fahlgren *et al*., 2015*b*). Based on the representative images, plants fluoresced more, retained less water, and produced less biomass when water was limited and salt was applied (Figures 5 and 8). This phenomenon was also detected by the FLUO and NIR cameras as shown in (Figure 6 and 10, Supplementary Figure 4 and 5). Plants under heat stress exhibited higher fluorescence under normal condition, but their water content did not change except for the day after the treatment was applied (Figure 15, Supplementary Figure 6). The heat conditions in the growth chamber led to a humidity increase (up to 85-90%) when temperature rose (data not shown), creating a positive environment for plant growth. When temperature increases, stomatal conductance increases, as well as transpiration and intercellular CO_2_ concentration (Urban *et al*., 2017). Our results for the heat experiment show a relatively lower NPQt (energy dissipated as heat) when plants were under stress conditions, indicating that the increase in humidity might contribute to the photosynthetic efficiency of the plants and relative water content (Figure 14). In agreement with this idea, the IR sensor had the ability to detect an increase in leaf temperature when the plants were under heat stress (Figure 13E-F). Higher temperatures promote flowering (Song *et al*., 2013). The IR sensor was able to detect differences in temperature across all genotypes at 23°C; the highest peak of temperature was reached twenty-four days after germination, which matched plant flowering time (Figure 13).

Fluorescence and near-infrared imaging reveal important information related to overall plant health, and when combined with the imaging power of the Scanalyzer HTS, it is possible to image multiple plants in rapid succession under identical conditions. Chlorophyll fluorescence directly relates to the rate of energy flow and electron transport within the plant, as well as the plant’s photosynthetic efficiency (Barbagallo *et al*., 2003; Murchie and Lawson, 2013). Over time, plants under normal water conditions had the tendency to emit increased fluorescent signal, which could be related to an increase in biomass and more available water (Figures 6C, 6F). On the other hand, when plants were subjected to 12.5% water limitation, they showed the largest area of “high” chlorophyll fluorescence compared with the control, which is considered an indicator of plant leaf senescence (Fahlgren *et al*., 2015*a*).

Another indicator of the effect of abiotic stress on plant physiology is yield. Plants under severe water limitation conditions produced the least seeds per plant for all genotypes (Figure 7). According to Basu *et al*., (2016), when growth conditions are extreme, plants cannot recover, and this may result in growth and/or yield penalty. When plants are exposed to a random short-term drought stress, they are able to mitigate the damage while they continue to grow and yield in challenging environments. However, a recent study has shown that when two stresses occur simultaneously the results are not always negative. Stress combinations can negate one another and potentially lead to a net neutral or even positive impact on plants (Pandey *et al*., 2017). Our data shows that when 50 mM NaCl was applied, soil water availability decreased (Figure 11), subjecting plants to a mild water limitation stress. This could explain why all genotypes (except *vtc*1-1) produced more seed under this salinity level, but as the level of salinity increased, yield dropped sharply (Figure 11).

### MIOX4 expression leads to abiotic stress tolerance

It has been shown that when over-expressing MIOX4, plants grew faster and accumulated more biomass than the controls (Lisko *et al*., 2013, 2014). For all extreme treatments (12.5%, 150 mM and 37°C), the restored lines accumulated more biomass than their respective controls (Figures 5C-D, 9C-D, 13C-D). According to Pavet *et al*., (2005), plants with low AsA content are more sensitive to heat and light stress compared with WT, and our data supports this statement. For example, when plants are subjected to water limitation, the stomata begin to close, lowering the CO_2_ uptake and limiting the Calvin cycle/increasing ROS in cells (Noctor *et al*., 2014). Plants under salt stress are altered nutritionally due to ion toxicity and osmotic stress. The increased Na^+^ concentration becomes toxic, which produces metabolic disorders and elevates ROS production (Acosta-Motos *et al*., 2017). Heat shock, on the other hand, induces oxidative stress in plants (Larkindale and Knight, 2002) and reduces AsA (Distéfano *et al*., 2017). As predicted from previous studies and mentioned before, there was a significant difference in all experiments when comparing abiotic stress tolerance. Under normal conditions, the restored lines accumulated more biomass, except at 23°C, where RV2 did not show a statistical difference but grew relatively larger than the rest of the genotypes.

It is possible that all the aforementioned responses correspond to the effect of over-expressing the MIOX4 ORF, which plays an important role in increasing AsA. This gene is also involved in the synthesis of cell wall polysaccharides (Kanter *et al*., 2005), which are considered protective barriers against abiotic stresses (Le Gall *et al*., 2015). Our data suggests that MIOX4 expression is higher when the restored lines are under abiotic stress, compared to no or very low expression of this gene in WT. This suggests that our restored lines and the MIOX4 line are producing more AsA due to the activation of the MI pathway (Figure 1B).

We hypothesize that MIOX4 plays a key role in AsA production during abiotic stress, which results in a significant increase in ROS scavenging to protect the plant from cellular damage. Additional work is necessary to identify the various components of this effect, especially assessments of expression with regard to other genes involved in the various AsA pathways as a proxy of the involvement of each branch in particular abiotic stresses. Exploring the subcellular distribution of AsA in these restored lines is also necessary. Subjecting lines to other stresses, such as biotic stresses, will assist in determining the effect of AsA under these circumstances. Such future studies should lead to a better understanding of the regulation of this important small molecule and its effect on abiotic stress tolerance.

## Supplementary data

**Supplementary Table 1.** Primers used for genotyping screening by PCR, sequencing and RT-qPCR.

**Supplementary Table 2.** Analysis of variance table from repeated measures analysis on the four responses, projected leaf area (cm^2^) and compactness, respectively.

**Supplementary Figure 1.** Seed germination per cross to determine homozygosis.

**Supplementary Figure 2.** Green (healthy) and yellow (chlorosis) expressed relative to projected leaf area as an indicator of tissue health in plants.

**Supplementary Figure 3.** Relationship between soil water saturation and plant water content/relative chlorophyll fluorescence.

**Supplementary Figure 4.** Representative images obtained using the Scanalyzer HTS (FLUO and NIR cameras) of Arabidopsis plants growing under 85% and 12.5% water saturation.

**Supplementary Figure 5.** Representative images obtained using the Scanalyzer HTS (FLUO and NIR cameras) of Arabidopsis plants growing under 0 mM NaCl and 150 mM NaCl treatment.

**Supplementary Figure 6.** Representative images obtained using the Scanalyzer HTS (FLUO and NIR cameras) of Arabidopsis plants growing under 23°C and 37°C.

## Acknowledgements

This research was funded by the Plant Imaging Consortium (NSF EPSCoR Track 2 Award #1430427) and the Wheat and Rice Center for Heat Resilience (NSF EPSCoR Track 2 Award #1736192). Authors thank support provided by the Arkansas Biosciences Institute and the Arkansas Research Alliance. We thank Dr. Fiona Goggin for facilitating collaboration between the Lorence and Lee groups. LMAG thanks the Molecular Biosciences PhD Program at A-State for partial stipend support.

## Authors constributions

Acosta-Gamboa LM designed, analyzed and performed the experiments. Acosta-Gamboa LM and Nepal N processed the experimental data, designed the figures and drafted the manuscript. Campbell Z and Cunningham SS processed experimental data. Medina-Jimenez K designed the primers used in order to detect the *vtc* mutations and aided in interpreting the results. Lee Jung Ae performed statistical analysis. Lorence A obtatined funding, design experiments, supervised the project and prepared the final version of the manuscript. All authors discussed the results and contributed to the final manuscript.

